# Brain size dependent speciation and extinction rates in birds and the cognitive buffer hypothesis

**DOI:** 10.1101/2025.02.04.634049

**Authors:** Jack W. Oyston, Michael R. May, Ryan N Felice

## Abstract

Both brain and body size vary greatly across birds, with both traits hypothesized to play a key role in diversification patterns across the group. Larger brains facilitate the evolution of greater behavioural plasticity, which could facilitate diversification in two ways, either by driving increased speciation (behavioural drive) or by making lineages more resilient to extinction (cognitive buffer). Here, we model brain size dependent diversification dynamics from published endocranial volume data for a sample of 2291 extant and fossil bird taxa, jointly modelling speciation rate, extinction rate and fossilisation rate separately for perching birds and other groups. We find that increasing relative brain size primarily increases diversification rate through decreasing extinction rate, supporting the cognitive buffer hypothesis and suggesting that larger brains in birds promote diversification by making species more resilient to extinction. Although speciation and extinction are dependent on relative brain size in both perching birds and other groups, this dependence is non-linear. In perching birds, the highest diversification rates are found over a narrow range of relative brain sizes, in contrast to other bird groups which show lower diversification rates over a broader range of relative brain sizes.

**Significance:** The potential for speciation and extinction are thought to be shaped by key phenotypic traits. In birds, brain size is one important trait hypothesized to have an impact on diversification. Testing this hypothesis requires the evolution of brain size, speciation rate, and extinction rate to be jointly modelled across the bird phylogeny. The inclusion of information on past diversity has been shown to be important in inferring diversification patterns, and so we include not only data from a broad sample of living birds but also information from the fossil record. Our study reveals patterns of brain size dependent diversification in birds which are consistent with the cognitive buffer hypothesis, with higher relative brain sizes being associated with lower extinction rates.

## Introduction

Patterns of speciation and extinction differ both within and between groups of organisms (1), leading to marked differences in taxonomic diversity across the tree of life (2). Understanding the factors driving speciation and extinction is key to understanding the origins of biodiversity, the macroevolutionary impact of external environmental forces, and the vulnerability of species to extinction. While it has been shown that climate and topography explain 60-80% of spatial unevenness of present-day global biodiversity in plant and vertebrate study systems (3), the mechanisms generating diversity over deep time are less well understood. Differences in the diversity of a clade through time result from the interaction of speciation and extinction with multiple factors such as climate change (4, 5), ecological niche availability (6) and population dynamics (7), as well as differences among organismal traits (8–10). The relative importance of these factors in driving the differential diversification of clades remains elusive in many cases (11), particularly for subclades within major groups which share the same bauplan and phenotypic traits and occupy similar ecological niches.

Birds are the most species-rich group of living terrestrial vertebrates and the only members of the Dinosauria to survive the Cretaceous-Paleogene (K-Pg) mass extinction 66 million years ago (Ma). Their long (ca. 165-150 Ma) evolutionary history (12) is punctuated by busts of rapid diversification (13–15), giving rise to a wide range of morphological and behavioural traits (16, 17) and making them ideally suited to studies of the macroevolutionary dynamics of speciation and extinction. Birds are ecologically varied and fulfil important roles in many ecosystems leading to them often being studied as indicator species (18–20). Many bird species are also at risk of extinction either currently or in the near future: around 23% of bird species are in IUCN Red List categories of concern (21). They have therefore been a clade of interest in diversification studies to inform conservation efforts in the context on the ongoing biodiversity crisis. Different groups of birds also show strikingly different diversity patterns. The perching birds (Passeriformes) are by far the most species-rich clade of birds, making up around 60% of extant bird diversity (22). The causes of this extreme unevenness in diversity have been much debated in the literature (23–25). Although both Passeriformes and non-passeriforms show size-dependent diversification trends, these trends cannot fully account for their differential species richness (26). Unlike other birds, high extinction rates are not associated with larger body sizes in Passeriformes (26), suggesting that other traits such as geographic isolation (27), sexual selection (28) or behavioural adaptations (29) likely contribute to the group’s exceptional diversity.

Numerous factors have been hypothesised to explain the evolution of bird diversity, but their relative importance continues to be debated. Despite higher diversity in the tropics, diversification rates show no trend with latitude (30). Instead, habitat age (31), precipitation (32) and vegetation (33) have been invoked to explain diversity differences between regions. While species richness within clades is expected to correlate with clade age (34), extrinsic factors such as density-dependent ecological limits (35, 36), range expansion (37) and migration (38) are also thought to have had a strong influence on the evolution of bird diversity. Organismal traits have also been proposed to shape patterns of species origination and extinction through time (39, 40). Among these traits, body size is often seen as one of the most important traits hypothesized to affect diversification in birds (41, 42). Speciation and extinction rates have been shown to be body size dependent in birds, with smaller bodied species diversifying many times faster than the largest-bodied flying lineages (26). Small body sizes might drive higher speciation rates through an association with higher mutation rates or shorter generation times (43). Although increased synonymous substitution rates are correlated with higher species richness in birds (44), these patterns could not be attributed to covariation with generation time or body size. Decreased diversification rates at larger body sizes might instead be associated with other life history traits, such as increased range sizes (45) or decreased population sizes (46). In addition to life history traits, other organismal traits are thought to affect species diversification, most notably brain size (43).

Several studies have shown that bird clades with larger average brain size tend to be more diverse (47, 48). In addition to changes in absolute trait values (body size, brain size), deviations from typical anatomical scaling relationships with body size brought about by changes in the underlying genetic and developmental regulatory mechanisms can also underpin major evolutionary shifts (49, 50).Such allometric changes can shape the trajectory of trait evolution on the macroevolutionary scale and underpins the origin of much present-day phenotypic diversity (51, 52). Brain size is often regarded as one of the most important allometrically varying traits and has been correlated with the evolution of important traits such as enhanced sensory capabilities (Walsh et al. 2018), cognition (53) and flight (54). Importantly, differences in relative brain size (correlated with cognitive ability) can be the result of changes in both body size and brain size (55). Although evidence suggests larger brains are associated with diversification in birds, the mechanism driving this association is more contentious. In the behavioural drive hypothesis, larger brains could facilitate the evolution of behaviours that expose individuals to new selective pressures and drive speciation (Wyles et al., 1983). In contrast, the cognitive buffer hypothesis (56, 57) predicts that larger relative brain size may also lead to reduced extinction rates, as larger brains allow enhanced learning capacity and greater behavioural flexibility which in turn allows them to better adapt to environmental changes and thus resistant to extinction threats.

The cognitive buffer hypothesis is broadly supported in ecological studies of both vertebrates (58–61) and invertebrates (62). It also seems to be supported in bird studies at the population level, as birds with larger relative brain sizes tend to have lower adult mortality rates (63), and reduced body size variation in response to climate warming (64). However, factors which may strongly affect brain size dependent diversification on microevolutionary scales might not hold true at macroevolutionary scales, particularly as relationships between brain size and body size become decoupled (65). Despite this, comparatively few studies have sought to test whether the cognitive buffer hypothesis holds true at macroevolutionary scales. A previous study, referred to hereafter as Sayol 2019 (66), found support for brain size dependent diversification patterns across the tree of extant birds. Sayol 2019 also found greatest support for a speciation-dependent model, over an equivalent extinction-dependent model or a null model with constant rates, leading them to conclude that brain size likely affected diversification rates by promoting speciation. This seems to suggest that while the cognitive buffer effect may be important over short time scales, the behavioural drive effect dominates at broader scales. While strong preliminary evidence that brain size does influence diversification in birds, the Sayol 2019 study has two main shortcomings in assessing the nature of these patterns. First, the models used did not allow both extinction and speciation rates to vary across the tree, likely due to difficulties in inferring extinction rates particularly without fossil taxa (67–69). We believe this is essential for accurately inferring whether brain size dependent diversification is driven by speciation, extinction, or an interaction of both. Second, the methods look at broad patterns across the entire bird tree. Songbirds are both highly diverse and have relatively large brains, but it remains unknown whether they show similar brain size dependent diversification patterns to birds as a whole. If different bird groups evolving large brains through different pathways, as seems to be the case in corvids and parrots (70), we might expect the impact of brain size on diversification dynamics to vary across groups. The few well-studied cases of adaptive radiations in birds are almost all perching bird groups, whether the Hawaiian honeycreepers (71), Madagascan vangas (72) or Galapagos finches (73), suggesting that at least certain passerine groups show different diversification patterns to those seen more broadly across birds. Testing whether large brains are associated with faster diversification rates in songbirds and whether these patterns are consistent with other bird groups is therefore instrumental in assessing the impact of brain size in shaping bird diversity patterns.

Different hypotheses regarding the effect of brain size on diversification rate make different specific predictions on how speciation and extinction rate should co-vary with relative brain size across clades. The state-dependent speciation and extinction (SSE) family of models (74, 75) are potentially highly useful for testing predictions of the effect of organismal traits on diversification, having been applied to investigating the effect of relative brain size (66), body size (76, 77), and flight ability (78, 79) in birds. These models have found consistent support for trait-dependent effects on diversification rate, but they have been less useful in distinguishing between underlying mechanisms, as this requires more detailed examination of the nature of these effects (beyond whether correlations are positive or negative) and whether they vary across groups. In addition, many studies of avian diversification do not include extinct taxa. It is known that incorporating information from the deep-time fossil record is necessary for reliably inferring diversification dynamics, particularly extinction rates (68, 69). While patterns of relative brain size have been investigated in birds in detail (70), this detail has not been reflected in corresponding models of diversification rate. If increases in relative brain size have shaped broad scale patterns of modern bird diversity (particularly high perching bird diversity), positive correlations between diversification rate and relative brain size should be recovered in other bird clades in addition to perching birds (Passeriformes). Such patterns being primarily the result of speciation or extinction rate would support either the behavioural drive or cognitive buffer hypothesis respectively.

There are strong theoretical and empirical reasons establishing a link between brain size and diversification rate in birds. In this study we seek to test whether this link is supported across the entire avian tree sampling both fossils and a broad subset of extant birds, as sampling the fossil record and accounting for fossilisation has been shown to impact the diversification patterns recovered. More importantly, we use this dataset to examine whether larger brains drive diversification primarily through increased speciation rate (‘behavioural drive’) or decreased extinction rate (‘cognitive buffer’) and whether these patterns could account for the exceptional diversity of the perching birds relative to other living bird groups.

Here we examine the effect of brain size on diversification in birds using published brain-volume data for a sample of 2291 extant and fossil birds, using Bayesian implementations of the quantitative state speciation and extinction (QuaSSE) family of models (75). Extensions to this approach (which we term BiQuaSSE models) allow estimation of transitions between different continuous trait dependent rate patterns for discrete traits or subgroups, essentially combining the structure of BiSSE and QuaSSE models. Further modifications for estimation of variable fossilisation rates and non-linear effects of traits on rate parameters allow us to test the predictions of the cognitive buffer hypothesis across the entire avian phylogeny.

Importantly the SSE models employed in this study are lineage-heterogenous not time-heterogenous. While time-heterogenous models have drawn criticism for issues with identifiability (69), lineage-heterogenous models are not susceptible to these issues (80). Trait-dependent lineage heterogenous models remain a strongly supported and statistically robust set of approaches which have been widely applied in evolutionary studies to provide insights into diversification and assess relative support for different models of evolution (81–83). In this study we quantify uncertainty in the parameter estimates of our QuaSSE models as well as evaluating their statistical power using simulated data (See Methods). We also compare our model results with a simulation based tip-rate method of testing for trait-dependent diversification (ES-sim)(84).

First, we test whether preferred models show net-diversification rate increases with increasing relative brain size in birds (HA1), or that net-diversification rate does not increase with increasing relative brain size in birds (HA0). Second, we investigate whether any such patterns are widespread across clades or limited to certain groups, such as the highly diverse songbirds. Specifically, we test whether differences in relative brain size track net-diversification rate in both the perching birds (Passeriformes) and other bird groups (HB1), or whether Passeriformes and non-Passeriformes show contrasting patterns for the effect of relative brain size on net-diversification rate (HB0). Third, we test the prediction of the cognitive buffer hypothesis that species with greater relative brain sizes are less likely to go extinct. If extinction rate decreases with increasing relative brain size across birds (HC1) this would support the cognitive buffer hypothesis. Conversely, if extinction rate does not decrease with increasing relative brain size across birds (HC0) then other mechanisms are likely to play a more important role in shaping macroevolutionary patterns of bird diversification, even if diversification rate increases with increasing relative brain size. Last, we test for evidence of a trend towards increasing relative brain size through time. Changes in relative brain size being biased in favour of increases rather than decreases (HD1) would indicate a trend towards increasing brain size throughout the evolutionary history of birds. If changes in relative brain size are not biased in favour of increases (HD0), this would indicate brain size either evolves randomly over long time scales, or that trends are local rather than across the entire bird phylogeny.

## Results

While extinction rates in birds are dependent on relative brain size (Fig. 1), speciation rates appear to be relatively constant. On average, speciation rates are only ∼4% higher in species with smaller relative brain sizes (Fig. 1A), with this decrease in speciation rates occurring gradually above relative brain sizes of -0.1. Birds with a relative brain sizes around -0.1 are mostly songbirds, including members of the thrush family (Turdidae) such as the fieldfare (*Turdus pilaris*) and tanagers (Thraupidae) like the blue-naped chlorophonia (*Chlorophonia cyanea*). Non-passeriformes with relative brain sizes around -0.1 include auks (Alcidae) like the whiskered auklet (*Aethia pygmaea*) and common guillemot (*Uria aalge*), and rails (Rallidae) like the South Island takahē (*Porphyrio hochstetteri*). Approximately 60% bird species in the dataset fall above this -0.1 threshold, including most songbird groups, corvids, parrots, penguins, birds of prey and woodpeckers.

**Figure 1:**
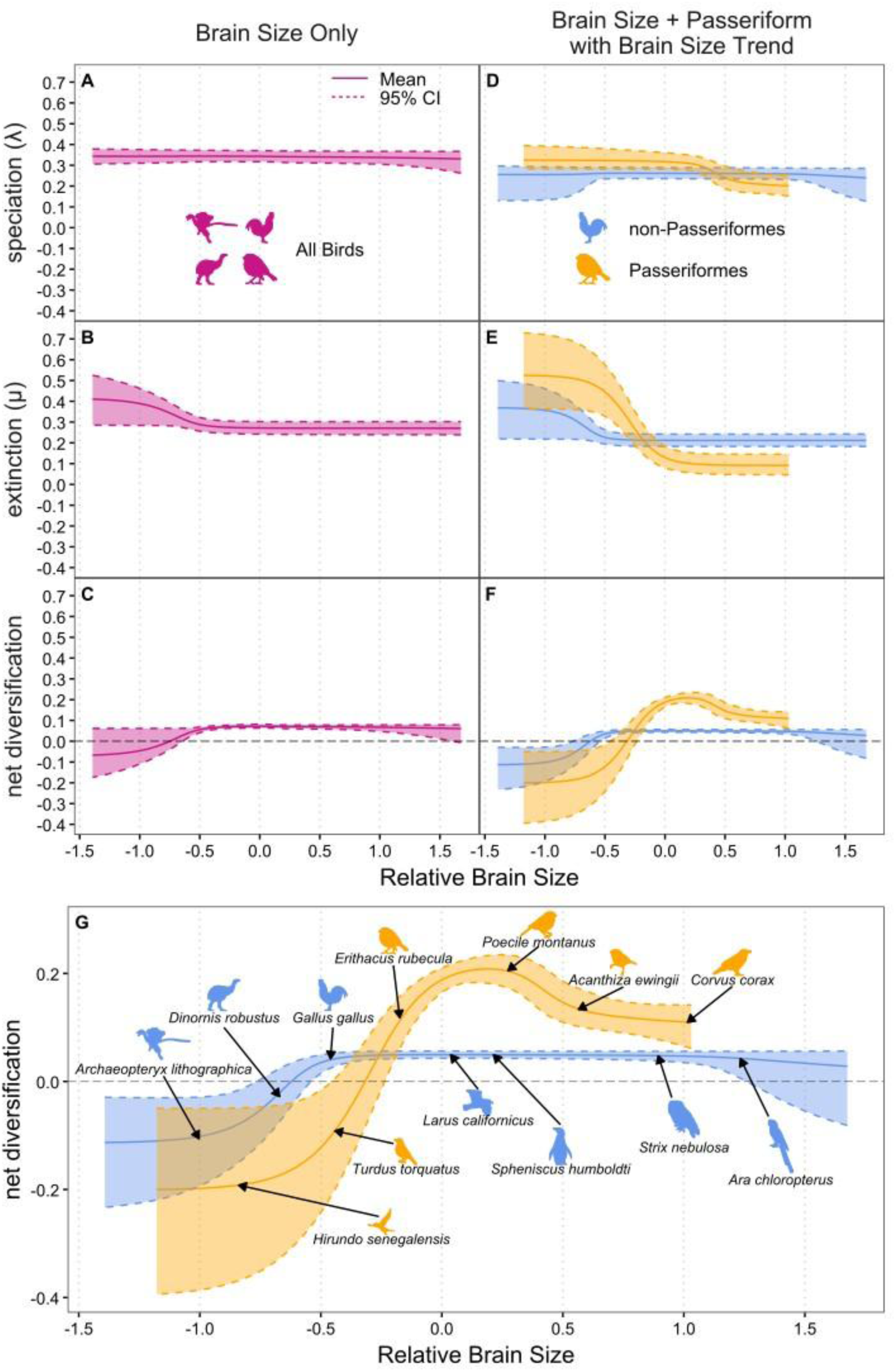
Diversification in birds (A-C) and for Passeriformes/non-Passeriformes (D-F) as a function of relative brain size. Residual brain size values from a linear regression against body size were used to control for the effect of allometry. Under the second model (D-F), the brain size transition rate is influenced by a bias term, allowing for trends in brain size evolution. Larger-brained lineages have lower speciation (A) and lower extinction (B) rates. Passeriformes have extinction rates lower than non-Passeriformes at highest brain sizes (E), leading to a peak in net diversification (F). Major groups of birds fall in different regions of the rate curves, with many songbird clades mapping onto the peak in net diversification (G). Model parameters inferred using the “middle crown” phylogeny (most recent common ancestor of crown birds at 92.51 Ma). See SI for results under alternative phylogenetic hypotheses and models. Animal silhouettes added to figures using the rphylopic R package (124).

Extinction rates vary slightly more with relative brain size than does speciation rate, birds with the smallest relative brain sizes having extinction rates ∼50% higher than birds with larger relative brains sizes (Fig. 1B). Extinction rates show a marked decrease at relative brain sizes between ∼-1.0 and -0.4. Around 14% of sampled bird species fall within the range of relative brain sizes associated with this decrease, including the oscillated turkey (*Meleagris ocellata*), common pigeon (*Columba livia*), great tinamou (*Tinamus major*), tawny pipit (*Anthus campestris*) and barn swallow (*Hirundo rustica*). Birds with relative brains sizes around -0.4 include the white wagtail (*Motacilla alba*), sharp-tailed sandpiper (*Calidris acuminata*) and the blue-crowned trogon (*Trogon curucui*).

Net-diversification rates in birds are positive and broadly show a slight increase with increasing relative brain size (Fig. 1C). The lowest net-diversification rates are shown by bird lineages with relative brain sizes below -1.0, the range of relative brain sizes associated with the highest speciation rates but also the highest extinction rates. These lineages include fossil stem birds, crown and stem Galloanserae (turkey, junglefowl, screamers) and paleognathes (rheas, moas and kiwis). Between relative brain sizes of ∼-1.0 and ∼-0.5 net-diversification rate more than doubles, due to declining extinction rates with increasing relative brain size as speciation rate remains relatively constant. Birds in this range include many Columbiformes (doves and pigeons), some Anseriformes (ducks), Coraciiformes (kingfishers, bee-eaters and rollers) and Podicipediformes (grebes). Net-diversification rates peak at relative brain sizes of -0.07-0.20. Lineages within this range include Piciformes (toucans and woodpeckers), Falconiformes (falcons and caracaras), Accipitriformes (vultures, eagles, kites and hawks), Fringillidae (true finches), Paridae (tits, titmice and chickadees) and Meliphagidae (honeyeaters). Above relative brain sizes of 0.20 there is a slight decline in diversification rates as speciation rates decline. Birds with relative brain sizes above 0.20 include bowerbirds (Ptilonorhynchidae), New World blackbirds (Icteridae), falcons (Falconidae) and penguins (Sphenisciformes). Net-diversification rates at the highest relative brain sizes are around 80% as fast as the peak net-diversification rates. Lineages with the highest relative brain sizes include some Strigiformes (owls), Cacatuidae and Psittacidae (cockatoos and holotropical parrots, respectively) and Corvidae (crow family). These patterns of net-diversification rate, as well as speciation and extinction, were similar regardless of differences in divergence time estimates (SI Figs. 1, 2 & 3).

Positive very weak correlations between relative brain size and diversification rate are also supported by alternative tests of trait based diversification in some cases, namely phylogenetic generalised linear models (PGLS) between equal splits measures of tip-based diversification and relative brain size (R2 = 0.002, p-value = 0.016, supplemental Table S1, figs. S10-12). Similar results are recovered for the extant only dataset (R2 = 0.002, p-value = 0.016, supplemental Table S1, figs. S10-12) but not for the passeriform of non-passeriform datasets individually. Simulation bases tests of equal-splits (ES-sim) were non-significant, yielding broad null distributions with many simulated correlation coefficients between -0.2 and 0.2.

Passeriformes (perching birds) and non-passeriforms show distinct relative brain size dependent diversification patterns when fitting BiQuaSSE models, which allow both groups to have different speciation and extinction rates (Fig. 1D-G). Passeriformes showing lower net-diversification rates than non-passeriforms at the lowest relative brain sizes, but markedly higher net-diversification rates at values above ∼-0.2 (Fig. 1F). At smaller relative brain sizes, speciation rates in Passeriformes are ∼25% higher on average than in non-passeriforms. In Passeriformes, a transition from higher to lower speciation rates occurs at relative brain sizes between 0.2 and 0.50 (Fig. 1D). Passeriforms falling within this range of decreasing speciation rates include species in Corvidae (the crow family), Ptilonorhynchidae (bowerbirds), tanagers (Thraupidae) and Cracticinae (currawongs & butcherbirds). Passeriformes with the lowest speciation rates are almost exclusively species in the genus *Corvus* (crows), with some bowerbird lineages (Ptilonorhynchidae) also showing low speciation rates. Non-passeriforms show much more constant speciation rates with changing relative brain size, although speciation rates are slightly lower at both the smallest and largest relative brain sizes. Speciation rates increase slightly with increasing relative brain size up to brain sizes of around -0.7, corresponding to many of the Galliformes, Columbiformes and paleognaths. The highest speciation rates in non-passeriforms are associated with relative brain sizes between -0.7 and 0.6. Waterfowl (Anseriformes) and shorebirds (Charadriiformes) fall on the lower end of this range, with Ibises, pelicans and birds of prey (Pelecaniformes and Accipitriformes) in the middle, and owls (Strigiformes) as well as hornbills and hoopoes (Bucerotiformes) at the upper end. The lowest speciation rates in non-passeriforms are associated with the largest relative brain sizes found largely within the parrots (Psittaciformes) as well as some true owls (Strigidae). Both groups also show higher extinction rates at lower relative brain sizes (Fig. 1E). In Passeriformes, the highest extinction rates are found in birds with relative brain sizes < - 0.40, including estrildid finches (Estrildidae), swallows (Hirundinidae) and tyrant flycatchers (Tyrannidae). Relative brain sizes between -0.40 and 0.09 show a rapid decrease in extinction rates to below those of non-passeriforms. Many songbirds, including most old-world flycatchers (Muscicapidae), thrushes (Turdidae) and Australian warblers (Acanthizidae) fall within this region. Passeriformes with relative brain sizes above 0.09 include wrens (Troglodytidae), cotingas (Cotingidae), crows (Corvidae) and bowerbirds (Ptilonorhynchidae). Non-passeriforms show highest extinction rates at relative brain sizes < -0.80, because of fossil taxa on the stem of major clades such as Aves (*Archaeopteryx*), Paleognathae (*Lithornis*) and Anserimorphae (*Presbyornis*). Between relative brain sizes of -0.81 and -0.30 extinction rates decrease. Birds with relative brain sizes within this range include many large ground birds such as Galliformes (gamefowl), Columbiformes (doves and pigeons) and paleognathes (particularly tinamous). Above relative brain sizes of -0.30 there is only a very slight and gradational decrease in extinction rate with increasing relative brain size in non-passeriforms.

The interaction between speciation and extinction rate ensures net-diversification patterns are dependent on relative brain size in both Passeriformes and non-passeriforms and are non-linear in both cases. Passeriformes show a pronounced peak in net-diversification rates with many songbird lineages showing positive values falling either on or directly either side of the maximum (Fig. 1G). Passeriformes and non-passeriforms show the lowest net-diversification rates in the smallest-brained taxa. Both groups then show an increase in net-diversification rate before a decrease in diversification rate at the highest relative brain sizes. The diversification rates of Passeriformes at the lowest relative brain sizes are roughly half (∼70% decrease) those of non-passeriforms with considerable uncertainty in net-diversification rates at these values, likely due few bird taxa having relative brain sizes in this range. There is then a rapid more than 4-fold increase in net-diversification rate between -0.6 and 0.2. Net-diversification rates then show a slight decrease in Passeriformes with relative brain sizes between 0.3 and 0.6. The increase in net-diversification rates with increasing relative brain size is more than 2-fold (140% increase) in non-passeriforms, occurring over a greater range of relative brain sizes (∼-0.8 to ∼-0.3). The highest net-diversification rates occur over a greater range of relative brain sizes (∼-0.3 to ∼0.7) in non-passeriforms, although diversification rates are much lower than in Passeriformes. Above relative brain sizes of ∼0.7, estimates of net-diversification rates in non-passeriforms are more uncertain, either remaining constant or decreasing slightly as relative brain size increases.

We recovered evidence that while relative brain size evolution is likely biased towards decreases in perching birds (Passeriformes), the distribution of biases in non-passeriforms is strongly impacted by the age of crown birds in the phylogeny (Fig. 2). When fitting QuaSSE models which allow asymmetrical transition rates between larger and smaller relative brain size categories, passeriforms show a broad negatively skewed distribution which is robust to uncertainty in the root node age of crown birds. In contrast, the distribution of the bias term in non-passeriform groups is only negatively skewed for the ‘middle’ age for the crown bird root node (92.51 Ma), being centered on 0 for the ‘early’ tree (110 Ma) and positively skewed for the ‘late’ tree (72 Ma). Decreases in relative brain size are therefore far more common than increases along Passeriform lineages, which we interpret as indicative of a trend towards decreasing brain sizes within the perching bird clade. We cannot conclude that the same is true in non-passeriforms as brain size changes are biased towards increases or decreases depending on the tree (SI Figs. 4, 5 & 6), but are more closely centered on 0 than those of passeriforms. The lack of a clear trend in relative brain sizes in non-passeriforms is reinforced by the absence of a clear bias in relative brain size evolution across the whole bird tree, as illustrated by ancestral state reconstructions of relative brain size according to a simple Brownian motion model (Fig. 3). Different bird clades within both Passeriformes and non-passeriforms show different trends towards increasing or decreasing relative brain size, again with many lineages showing a trend towards decreasing relative brain size.

**Figure 2:**
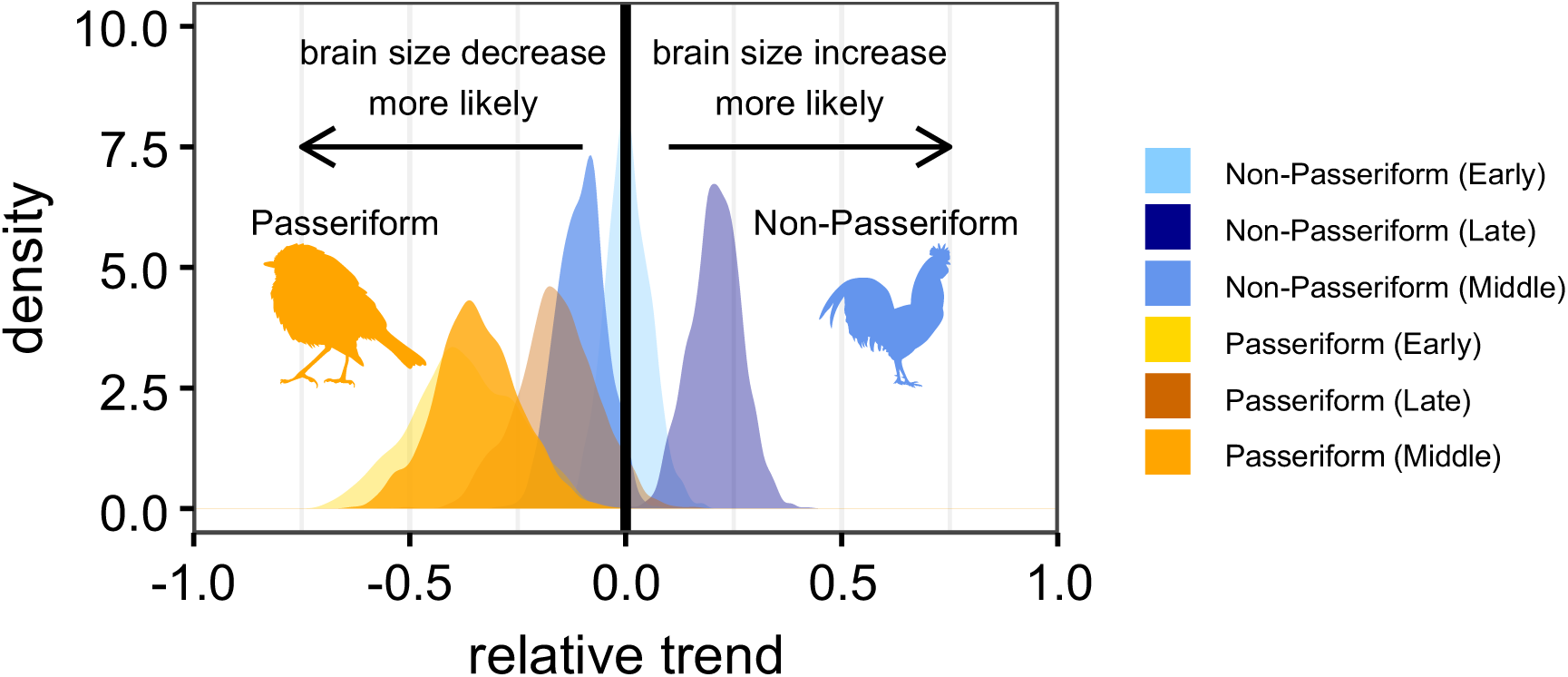
Estimated trends in relative brain size transitions. Values greater than zero indicate a bias toward brain size increases, values lower than zero indicate a bias toward brain size decreases. Both Passeriformes and non-Passeriformes tend towards decreases in brain size more often than increases, although this trend is relatively weak in non-Passeriformes. The “Early”, “Middle” and “Late” phylogenies set the age of the most recent common ancestor of crown birds at 110 Ma, 92.51 Ma and 72 Ma respectively.

**Figure 3:**
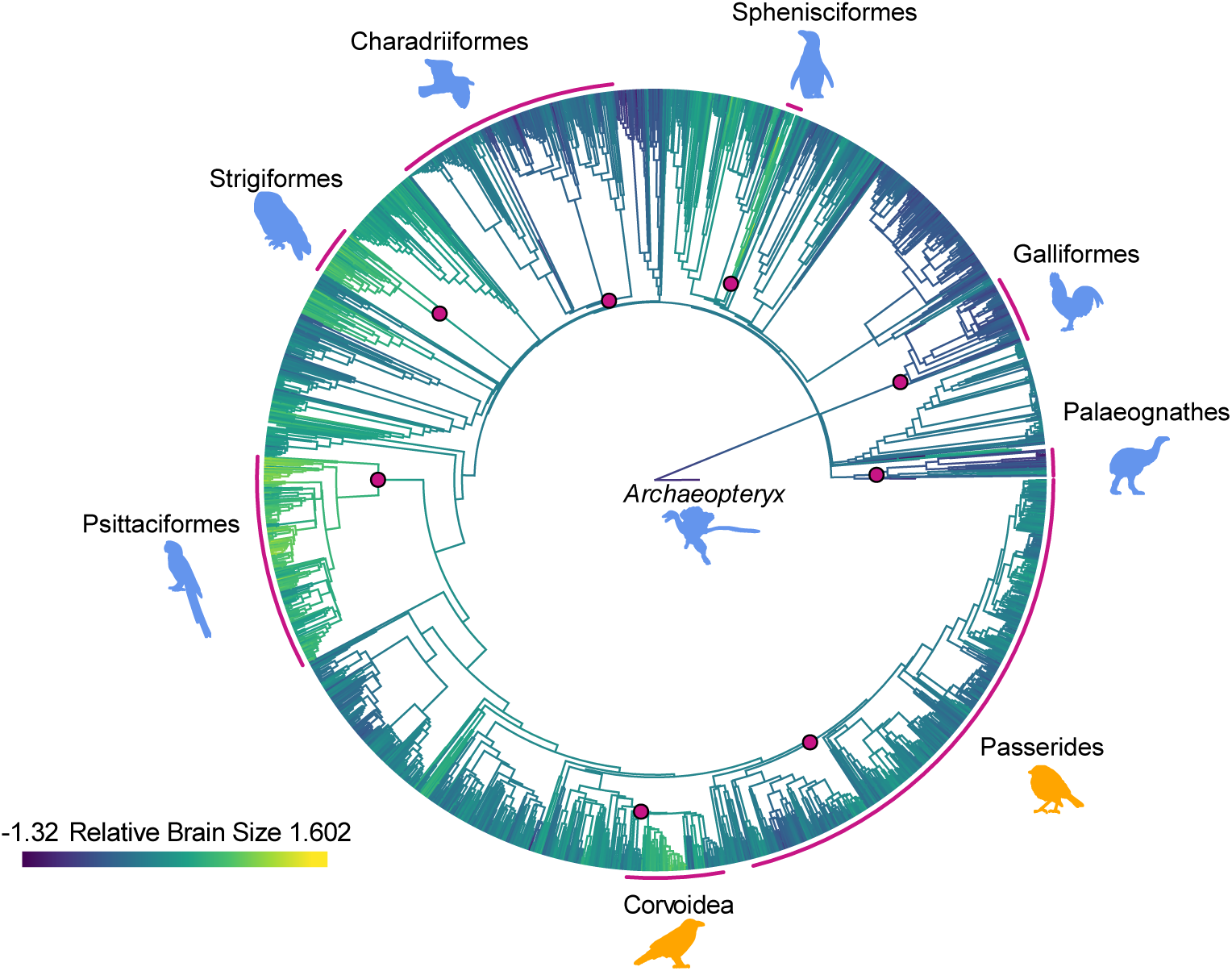
Evolution of bird relative brain size over time. Relative brain sizes were estimated and plotted in the R package phytools (125). Ancestral states were inferred using fastanc under a simple Brownian motion model. Traits were then mapped to the phylogeny using the contMap function.

## Discussion

Our analyses add to the growing body of evidence for brain size influencing diversification rate, supporting previous findings that bird lineages with larger relative brain sizes consistently exhibit higher net-diversification rates (66). However, our results differ from previous findings in that we found that this increase in diversification rate at larger relative brain sizes was driven primarily by decreasing extinction rates, rather than increasing speciation rates. Our results are consistent with the “cognitive buffer hypothesis”, namely that increasing relative brain size promotes diversification, through facilitating the development of plastic behavioural responses. Greater behavioural plasticity likely allows those taxa with larger brains to modify their behaviour to better respond to environmental changes and resource shortages, making them more resilient to extinction. Furthermore, the fact that both Passeriformes and non-passeriforms show this pattern of lowest diversification rates and highest extinction rates at low relative brain sizes suggests that these trends correspond to changes in relative brain size rather than being unique to Passeriformes.

Although PGLS-based tests support a significant correlation between tip-rate and relative brain size, these correlations were both extremely weak and not supported by complementary simulation-based methods (ES-sim). This lack of support for strong linear correlations between relative brain size and diversification is consistent with the non-linear relationships supported by QuaSSE models; across all birds net diversification rate increases only slightly with increasing relative brain size. Additionally, although used as a metric of diversification, tip-rate correlation methods are based on the times between branching events and are therefore likely to be more reflective of speciation rate rather than extinction rate (84, 85). As speciation rate appears independent of relative brain size across all birds it is unsurprising that inverse equal splits measures which capture variation in branching rate, a proxy for speciation rate, is only very weakly correlated or uncorrelated with relative brain size.

This divergence from previous inferences about relative brain size dependent diversification is somewhat surprising, but our analyses differ from those of the Sayol 2019 in several aspects, both in terms of the diversification rate models used and the underlying dataset. Our dataset is larger than those of previous studies (n = 2291 compared to n = 1931, a difference of 18.6%). Most importantly, to our knowledge this is the first study of diversification rate in birds to include estimates of brain size from extinct taxa. Including fossil stem bird taxa and early representatives of clades within Neoaves notably increases the range of brain size relative to body size. Basally diverging crown birds show a weaker scaling relationship between brain and body size, with rapid shifts in relative brain size occurring in the aftermath of the K-Pg extinction (70). Accordingly, taxon sampling, rather than differences in topology or specimen sampling, had the biggest impact on our estimates of relative brain size. We carried out sensitivity analyses on the dataset with fossil taxa removed and recovered a slight increase in speciation rate with increasing relative brain size (SI Figs. 10, 11 & 12), broadly consistent with the results of previous studies (66).

The QuaSSE models used by Sayol 2019 modelled speciation and extinction rates as a linear function of relative brain size. Our results support a non-linear relationship between diversification rate and relative brain size in birds, even accounting for uncertainty in rate estimates at low relative brain sizes in the fitted sigmoid functions. Both speciation and extinction rates in the reduced extant-only dataset have higher associated error and a lower effective sample size. This is particularly notable at lower relative brain sizes, likely due to many fossil birds being large with relatively small brains. Furthermore, these previous QuaSSE analyses (Sayol 2019) also kept either speciation rate or extinction rate independent of relative brain size (the same rate across all trait values). As net-diversification was found to increase with increasing relative brain size in all cases, this would necessarily support increasing speciation rates if extinction rate were fixed. Although the model with a fixed extinction rate was preferred over the null constant model or the model with a fixed speciation rate, this could be biased by underestimates in extinction rate when analysing only extant taxa. In our analyses we allowed both speciation rate and extinction rate to vary with relative brain size, constrained by the estimated sampling of the bird phylogeny, dates of the tips in the tree (notably fossils), and the fossilisation rate. In both Sayol 2019 and this study relative brain size dependent trends in speciation rate are modest. It is therefore not surprising that here was uncertainty in our QuaSSE models estimations of the direction of the slope for speciation rate. The interaction of speciation and extinction rate contributing to net-diversification meant that some models with modest positive slopes for speciation rate (increasing speciation rate with increasing relative brain size) were about as likely as models with modest negative slopes for speciation rate (decreasing speciation rate with increasing relative brain size), depending on the strength of the estimated negative slope for extinction rate (how steeply extinction rate decreases with relative brain size and the shape of the sigmoid curve). Taken together, the larger taxon sample, the inclusion of fossil taxa and not assuming a linear relationship between rate parameters and relative brain size allows the present study to more accurately infer trait-dependent diversification patterns, particularly regarding speciation rate.

There are reasons to expect greater behavioural flexibility might not promote more rapid speciation. If large-brained species can use plastic behavioural responses to move close to adaptive optima sooner than they would through phenotypic evolution, then this should mask genetic variation from natural selection as the selection on the phenotype is reduced (86, 87). Indeed, there is evidence that larger-brained species have longer generation times and slower life history strategies in general (48, 57), due to the high energetic cost of developing large brains lengthening development time. This in turn would reduce the frequency of allelic changes and likely the per time mutation rate along branches (88), at least for neutral mutations. Depending on the strength of selection, this effect might be expected to reduce speciation rates. In contrast a link between larger brains and decreased extinction rates seems theoretically clearer, as greater behavioural flexibility allows species to adapt more quickly to changing environments and resources (89, 90). There is also evidence that large brains make taxa better able to expand their ranges (91) and establish themselves in new areas (92–94). While this could theoretically promote allopatric speciation, there is no evidence that birds with larger brains are better able to colonise remote areas such as volcanic islands (95). It does suggest, however, that larger brains in birds does translate into traits which are likely to make species more resilient to extinction.

Relative brain size dependent diversification patterns are non-linear, with birds showing decreased diversification rates in the largest-brained lineages due to declining speciation rates. This peak is most pronounced in the perching birds (Passeriformes), although other bird groups show evidence of a broader, flatter peak indicating that this pattern holds true for birds generally. This suggests that there is a trade-off between a benefit of larger brains in accordance with the cognitive buffer hypothesis and other factors which confer an evolutionary cost to larger brains. Our results would indicate that there is an optimal range of relative brain sizes in birds associated with the highest diversification rates which is most clearly seen when looking at the effect of relative brain size on net-diversification rate across all birds. Most obviously, the high energetic costs associated with developing larger brains might impose constraints on brain size evolution which select against larger brains above a threshold value. As discussed previously, larger brain size is also associated with longer generation times and slower mutation rates (48, 57) which could lead to lower speciation rates. If the evolution of large brains above a certain threshold only occurs when there are strong evolutionary drivers to shift the balance of these trade-offs, perhaps driven by strong selection for behavioural flexibility in order to exploit specific resources or as an adaptation to niche, highly variable environments, then the largest brains could represent a form of phenotypic specialisation which limits the ability of those lineages to subsequently diversify. Both parrots (Psittaciformes) and the crow family (Corvidae) have relatively low species richness, despite having the highest average relative brain sizes within birds. In both groups large brains are associated with enhanced cognitive abilities and associated morphological traits specialised to manipulate objects to take advantage of hard-to-access food resources. Although this study focused on relative brain size as a measure which is independent of the allometric relationship between brain and body size, body size could still influence brain size trends indirectly. Both speciation and extinction rates are known to be strongly body size dependent in birds, with smaller birds having higher diversification rates (26). The scaling relationship between brain size and body size is likely weakly curvilinear (96), so linear approximations might tend to underestimate relative brain size at larger body sizes. In addition, other traits such as range size (97), habitat variability (98), nest structure (99) and eye size (100, 101) likely covary with both brain size and body size. Investigating the influence and interaction of these different traits on diversification patterns is beyond the scope of this study but represents a much needed and likely fruitful avenue of future research in the field.

Perching birds (Passeriformes) show stronger brain size dependant effects than other birds despite exhibiting a smaller range of relative brain sizes, with both higher and lower speciation and extinction rates than other groups. Most noticeably, they show a stronger non-linear pattern than non-passeriforms with a sharp peak in net-diversification rates at relatively large brain sizes corresponding to many songbird lineages. The sigmoid curves describing the influence of brain size on speciation and extinction rates are also offset compared to non-passeriforms, showing a sharper decrease in speciation rate at lower relative brain sizes than non-passeriforms but also a much sharper decrease in extinction rates at higher relative brain sizes than non-passeriforms (Fig 1D-E). These two patterns combine to produce the sharp peak in Passeriform diversification rates associated with a narrow range of relative brain sizes when compared to other groups, with most of the hyper-diverse songbird clades corresponding to relative brain sizes either at or slightly smaller than this peak (Fig 1F-G). Conversely, other groups show a flatter peak of relative brain sizes with lower diversification rates. This would indicate that while relative brain size influences speciation and extinction rate in both groups the trade-offs between the benefits and disadvantages of increasing brain size on diversification are different for Passeriformes, such that the high diversity of perching bird groups cannot be explained solely as a function of increased relative brain size. Other explanations such as greater habitat variability (102), more frequent geographic isolation of Passeriform lineages (103, 104), or the evolution of complex songs (105) leading to reproductive barriers, but see (106, 107), may explain why Passeriformes depart from the patterns seen in other bird clades. It is possible that the cognitive buffer effect in perching birds is reduced compared to lineages from other groups with the same relative brain size due to sexual selection for large brains in perching birds to produce complex songs (108, 109) rather than for greater behavioural plasticity. Although there are both passeriform and non-passeriform groups with exceptionally large relative brain sizes (Corvidae and Psittaciformes respectively), crows and parrots appear to have evolved large relative brain sizes through markedly different routes (70). While parrots evolved large relative brain sizes primarily through decreases in body size, corvids evolved increased body size while increasing relative brain size at a faster rate. The sharper decrease in speciation rates in Passeriformes could be a consequence of evolving larger body sizes in Corvidae in association with larger brains in Corvidae but not Psittaciformes, as speciation rate tends to decrease with increasing body size (26).

Contrary to our expectations and despite increased diversification rates at larger relative brain sizes, we recover support for a bias towards decreases in relative brain size in evolving perching bird lineages. This contrasts with trends recovered for body size which show a strong widespread bias towards increases (26). The distribution of increases and decreases in non-passeriforms was dependent on the age used for the root node of crown birds, being more biased towards increases in relative brain size when a later age for crown birds is used. This would imply that large increases in relative brain size tend to occur on basal branches within major clades of birds. If a later age is used for the last common ancestor of crown birds, these basal branches tend to be shorter, resulting in the same increases in relative brain size on these branches occurring over shorter periods of time. This will, in turn, favour positive bias terms (increases more likely than decreases). This observation is consistent with reported patterns of brain size evolution across birds, namely rapid increases in relative brain size early in Neoaves followed by contrasting trends in different major clades of extant birds subsequently (70). Different evolutionary trajectories towards clade-specific optima in modern Neoaves would also explain why we failed to recover evidence for trends in relative brain size more broadly across birds. Most of the relative brain size transitions in Passeriformes are negative, suggesting that there is a bias towards decreasing relative brain sizes in this group. The fact that many passerine clades have small body sizes coupled with the bias towards increasing body size in bird lineages suggests this bias towards decreasing relative brain sizes may be due to a tendency to evolve larger body sizes without corresponding increases in brain size. Many perching birds may be constrained at or close to their minimum body sizes such that further decreases in body size are not selected for. Constraints operating at small body size in perching birds include morphological constraint of vocalisations, particularly bandwidth (110) and thermal constraints corresponding to Bergmann’s Rule (111). This may favour decreases in relative brain size if selection for increased body size is stronger than selection for increased brain size.

Through the application of trait dependent diversification models which jointly model both speciation and extinction rate non-linearly, we have revealed novel insights into the importance of relative brain size on diversification dynamics in birds. Diversification is brain size dependent in birds in a non-linear fashion, with narrow range of relative brain sizes associated with the highest diversification rates. By incorporating fossil taxa and increasing the taxon sample size we can demonstrate directly for the first time that brain size has a pronounced effect on extinction rate in birds, providing compelling support for the cognitive buffer hypothesis. While both perching birds (Passeriformes) and other groups (non-passeriforms) show brain size dependent diversification processes, differing patterns within the two groups are likely the result of differences in the trade-offs between reduced extinction rates and reduced speciation rates at larger relative brain sizes. Passeriformes show markedly lower extinction rates, suggesting that while high diversity in perching birds is partially result of the evolution of large brains allowing passerine lineages to persist for longer. Taken together, these results highlight the complex effect of brain size evolution on diversification patterns within birds as well as differences in the interplay between brain size and body size in different bird clades.

## Methods

### Brain volume data

Brain volume estimates of birds (Avialae) were compiled from the recently published literature. Estimates for 305 extinct and living species were obtained from the CT-rendered endocast dataset of Ksepka et al. 2020, lead shot braincase volume measurements of 1931 extant species taken from Sayol 2019 and estimates for 360 extant and fossil species taken from papers published after 2020 (See Auxillary SI). For two taxa (unidentified galliform SMF_Av_666 (112) and *Spheniscus urbinai* (113)) endocast volumes were estimated directly from CT-scan 3D surfaces, in all other cases volume estimates were taken from the published values given in the source publications referred to above. In cases where there were multiple entries with different values for the same species (such as when different specimens were measured or averaged in different studies), the mean of these entries was taken as the value for that species. This resulted in a dataset of 2291 bird species (2260 extant and 31 extinct).

### Body size data

Body size data for extant taxa were taken from the AVONET database (114), while values for extinct taxa were taken from the dataset of Felice et al. (26). 8 species included in this study were missing from this dataset (unidentified galliform SMF_Av_666, unidentified *Presbyornis*, *Procariama simplex*, *Spheniscus urbinai*, *Otus murivorus*, *Scandiavis mikkelseni*, *Streptopelia risoria* and *Centuriavis lioae*). In 7 cases body size estimates were taken directly from published values in the source publications (See Auxillary SI), in 1 case (*Spheniscus urbinai*) body size estimates were calculated from linear measurements of long bones using established scaling equations (115).

### Phylogenetic Hypotheses

Phylogenetic trees were based on the informal supertree of Felice et al. (26), based on the Hackett backbone topology grafted to the recent Yu et al. phylogeny (116) focused on Cretaceous birds. The ‘tree.merger’ function of the R package Rrphylo (117) was used to graft the 6 taxa not already included in this phylogeny for which estimates of brain volume were available onto the composite tree. Phylogenetic relationships and tip dates for these 6 taxa were derived from the primary literature (See Auxillary SI). In cases where fossil ages were provided as subseries or stages the ICS Chronostratigraphic Chart 2024 (118) was used to assign absolute ages. Uncertainty in the origination of the crown group was accounted for by rescaling branches in the tree using the ‘scaleTree’ function of the RRphylo package to modify the age of the most recent common ancestor of *Struthio camelus* and *Passer domesticus*. These modified ages matched the alternatives used in Felice et al. (26): “early crown” (110 Ma), “middle crown” (92.51 Ma) or “late crown” (72 Ma). Lastly, the tree was pruned down to include only the 2,091 species included in the brain volume dataset (See Auxillary SI) using the keep.tip function in the R package ape (119).

### Models of continuous trait evolution

Brain size scales allometrically as a function of body size. Additionally, patterns of brain size across the bird phylogeny are also in part due to shared ancestry (phylogenetic relatedness) which could confound relationships of how brain and body size covary. Residuals of a phylogenetic linear regression of log body size (x) against log brain volume (y) using the R package phylolm were therefore the preferred metric of relative brain size for this study, as this measure of relative brain size takes into account both allometry and phylogeny. For each metric, the mean value was then subtracted from the value for each species ensuring both distributions had a mean of 0 prior to analysis. We fit relaxed phylogenetic Brownian motion models of continuous trait evolution in RevBayes (120) using the function dnPhyloBrownianREML. We fit three models to each measure of brain size: relaxed Brownian motion, Brownian motion with a trend and Brownian motion with a trend and tips partitioned such that passeriform and non-passeriform lineages may have different rate and trend parameters (121). We also fit multivariate equivalents of these same three models to both relative brain volume and log body size data. For each analysis we ran 12 chains of 10,000 generations with 16 moves per generation, discarding the first 1000 generations as burnin and tuning prior distributions every 5 generations.

### Trait-dependent diversification models

Relative brain volume dependent speciation and extinction models, as well as multivariate models dependent on both relative brain volume and log body mass were implemented in RevBayes using custom code. First, continuous traits were discretised into 32 bins using a custom script in RevBayes. These data were then input into QuaSSE models of trait dependent diversification in RevBayes. Sigmoid functions to model the effect of relative brain volume and body size on speciation rate, extinction rate and mass extinction probability, while fossilisation and sampling rate were kept constant through time and across all trait bins. We fit two Brownian motion models simulating relative brain size evolution in the QuaSSE models, one where increases and decreases in relative brain size were symmetrical (SI Figs. 7, 8 & 9), and one specifying separate rate parameters (sigma up/ sigma down) for transitions to larger or smaller relative brain size categories respectively. Each model consisted of 12 MCMC chains run for 10,000 generations with 23 moves per generation and tuning prior distributions every 5 generations for the first 1,000. The first 20% of each chain was removed as burn-in and any runs that failed to achieve suitable ESS were rejected. In addition to these QuaSSE models, trait dependent diversification models where tips were partitioned such that passeriform and non-passeriform lineages may have different rate and trend parameters were also fitted to the data (BiQuaSSE models). In the case of the BiQuaSSE models, the discrete categories of the continuous traits were then combined with the binary codings for the discrete trait of interest (whether the taxon was passeriform or non-passeriform) using a custom R script. This transformed the columns of continuous trait values and the column of discrete trait codings into a single column of 64 categorical bins (1-32 as codings for the bins of non-passeriform and 33-64 as codings for the same bins for passeriforms). BiQuaSSE models used separate sigmoid functions for passeriforms and non-passeriforms to model the effect of relative brain volume and body size on speciation rate, extinction rate and fossilization rate. Mass extinction probability was allowed to vary also for passeriform and non-passeriform birds but was kept constant within each category, while sampling was kept constant through time. Again, for the BiQuaSSE analyses we fit two models simulating trait evolution under a Brownian motion model and Brownian motion allowing different rates for transitions between larger and smaller relative brain size categories. In all BiQuaSSE models model parameters were allowed to differ between passeriforms and non-passeriforms. Each model consisted of 12 MCMC chains run for 10,000 generations with 24 moves per generation, discarding the first 2500 generations as burnin and tuning prior distributions every 5 generations for the first 1,000 generations. The first 20% of each chain was removed as burn-in and any runs that failed to achieve suitable ESS were rejected.

### Sensitivity Analyses & Simulations

Correlations between tip rate measures of diversification using the equal splits method (84) and traits show lower false discovery rates than QuaSSE analyses, which can be prone to false positives in some cases (122). Equal splits uses the log-transformed inverse equal splits (ES) measure to quantify diversification at each tip as the sum of edge lengths subtending it. Significance was then evaluated using a simulation test of 1,000 replicates of Brownian trait evolution across the tree (ES-sim), as well as phylogenetically corrected linear models where ES was dependent on relative brain size given the tree using the pgls function in caper (123). ES-sim tests (SI Tables 1 & 2) were performed on different data partitions (fossil/extant and passeriform/non-passeriform) as well as on the 3 phylogenies corresponding to different ages for the root node of crown birds (early, middle and late). ES-sim analyses were carried out in R using the code available from the Harvey Lab github https://github.com/mgharvey/ES-sim (See Auxillary Supplemental Data scripts).

The statistical power of trait-dependent diversification models was evaluated using simulated data. Phylogenetic trees and relative brain size data were simultaneously forward simulated using the sigmoid curves and input parameters derived from the posterior mean estimates of our fitted empirical models. The trait dependent diversification parameters were then estimated using the same methods applied to estimating values for the empirical data. All models were fit with 12 MCMC chains of 10,000 generations each. The first 20% of each chain was removed as burn-in and any runs that failed to achieve suitable ESS were rejected. The estimates of parameter values from these runs were evaluated using 3 metrics: the coverage (frequency that the simulated true value falls in the 95% confidence interval of the posterior distribution of estimated values), precision (the percent difference between the simulated true value and posterior mean estimate of the value), and accuracy (residual mean squared error).

## Notes

### Competing Interest Statement

The authors have declared no competing interest.

