## Supplementary material for "Brain size dependent speciation and extinction rates in birds and the cognitive buffer hypothesis": SI

Supplementary Materials

QuaSSE Model Results

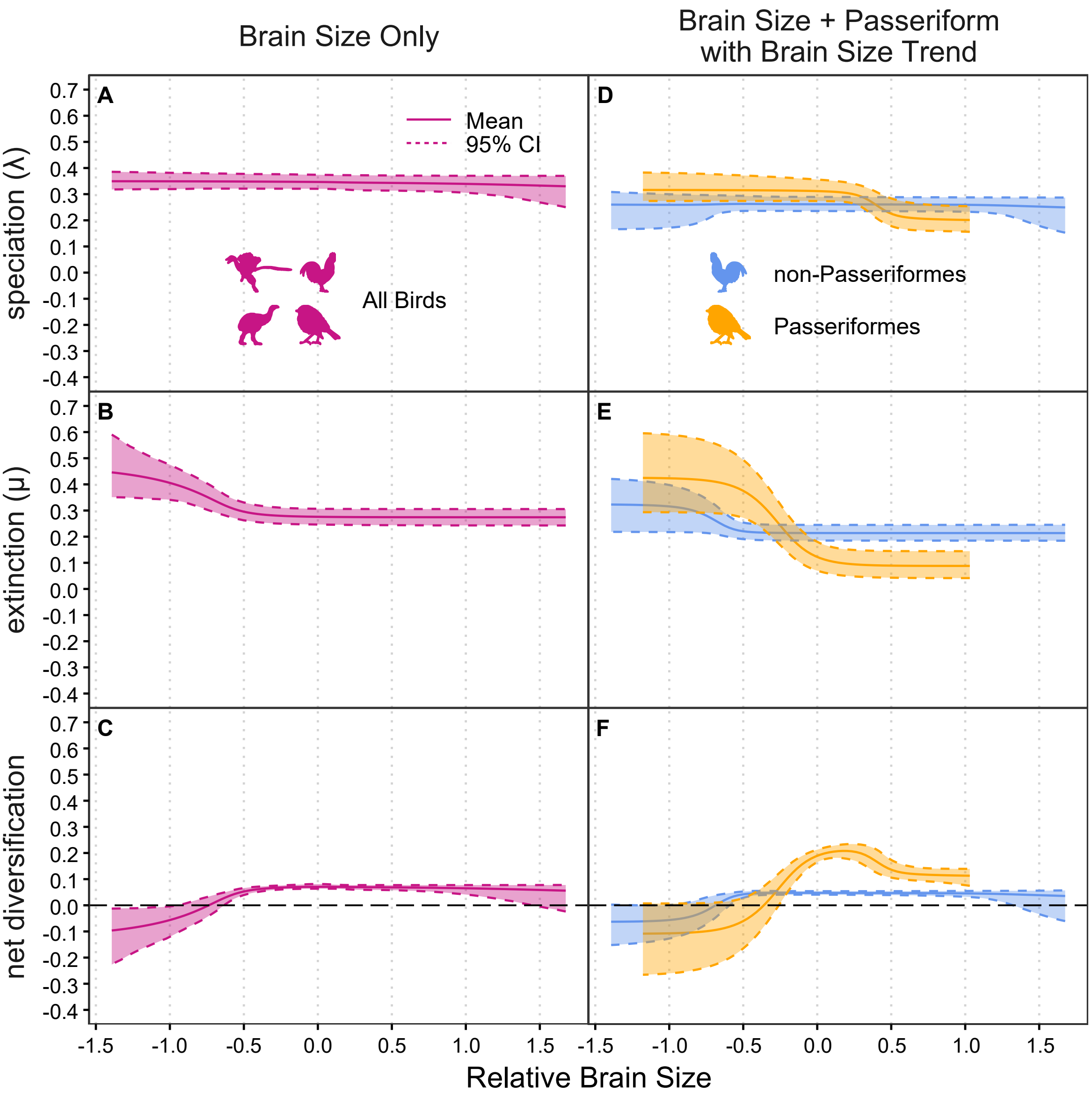


Fig 1: Diversification in birds (A-C) and for Passeriformes/non-Passeriformes (D-F) as a function of relative brain size. Residual brain size values from a linear regression against body size were used to control for the effect of allometry. Under the second model (D-F), the brain size transition rate is influenced by a bias term, allowing for trends in brain size evolution. Larger-brained lineages have lower speciation (A) and lower extinction (B) rates. Passeriformes have extinction rates lower than non-Passeriformes at highest brain sizes (E), leading to a peak in net diversification (F). Model parameters inferred using the “early crown” phylogeny (most recent common ancestor of crown birds at 110 Ma).

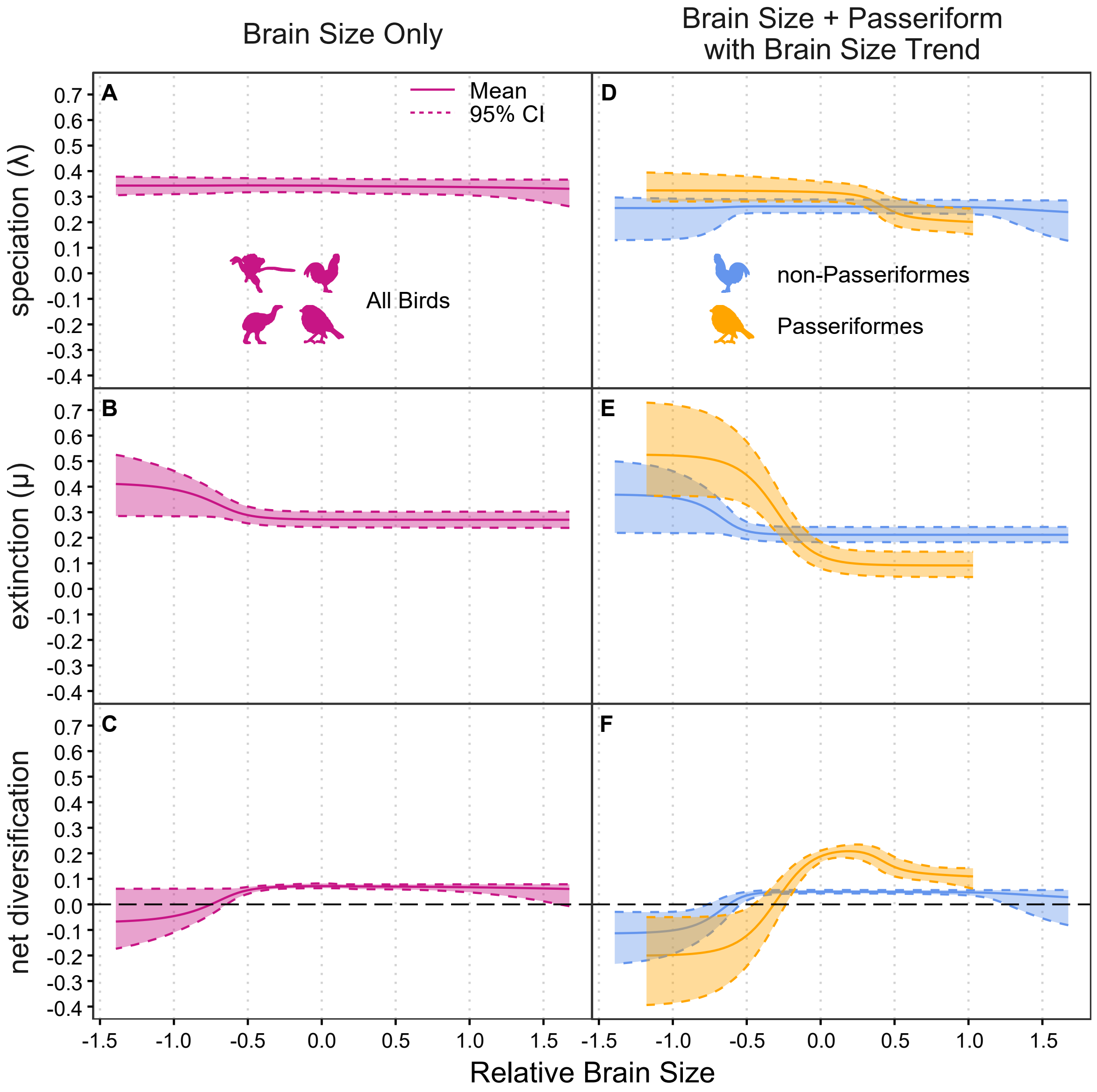


Fig 2: Diversification in birds (A-C) and for Passeriformes/non-Passeriformes (D-F) as a function of relative brain size. Residual brain size values from a linear regression against body size were used to control for the effect of allometry. Under the second model (D-F), the brain size transition rate is influenced by a bias term, allowing for trends in brain size evolution. Larger-brained lineages have lower speciation (A) and lower extinction (B) rates. Passeriformes have extinction rates lower than non-Passeriformes at highest brain sizes (E), leading to a peak in net diversification (F). Model parameters inferred using the “middle crown” phylogeny (most recent common ancestor of crown birds at 92.51 Ma).

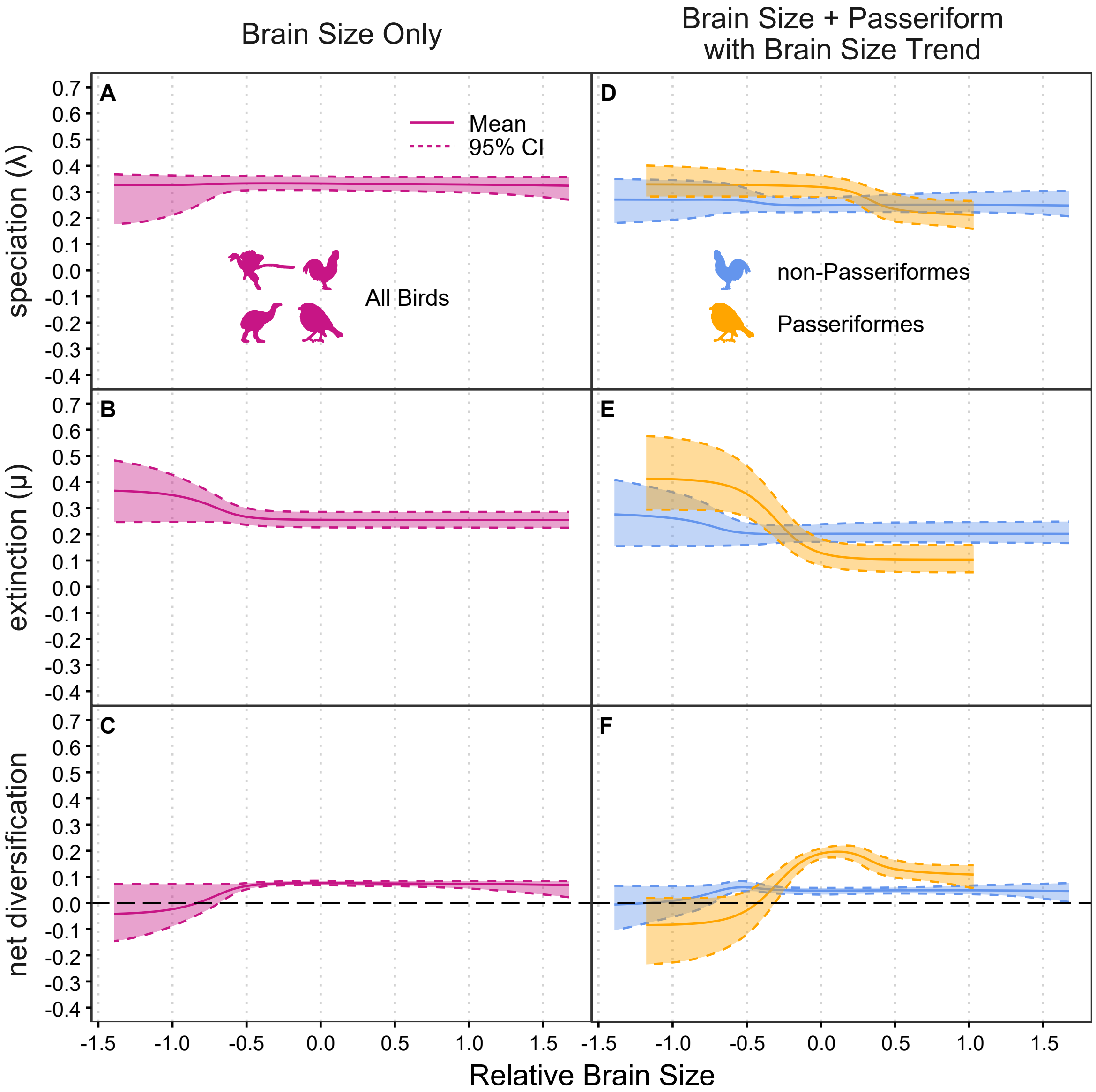


Fig 3: Diversification in birds (A-C) and for Passeriformes/non-Passeriformes (D-F) as a function of relative brain size. Residual brain size values from a linear regression against body size were used to control for the effect of allometry. Under the second model (D-F), the brain size transition rate is influenced by a bias term, allowing for trends in brain size evolution. Larger-brained lineages have lower speciation (A) and lower extinction (B) rates. Passeriformes have extinction rates lower than non-Passeriformes at highest brain sizes (E), leading to a peak in net diversification (F). Model parameters inferred using the "late crown" phylogeny (most recent common ancestor of crown birds at 72 Ma).

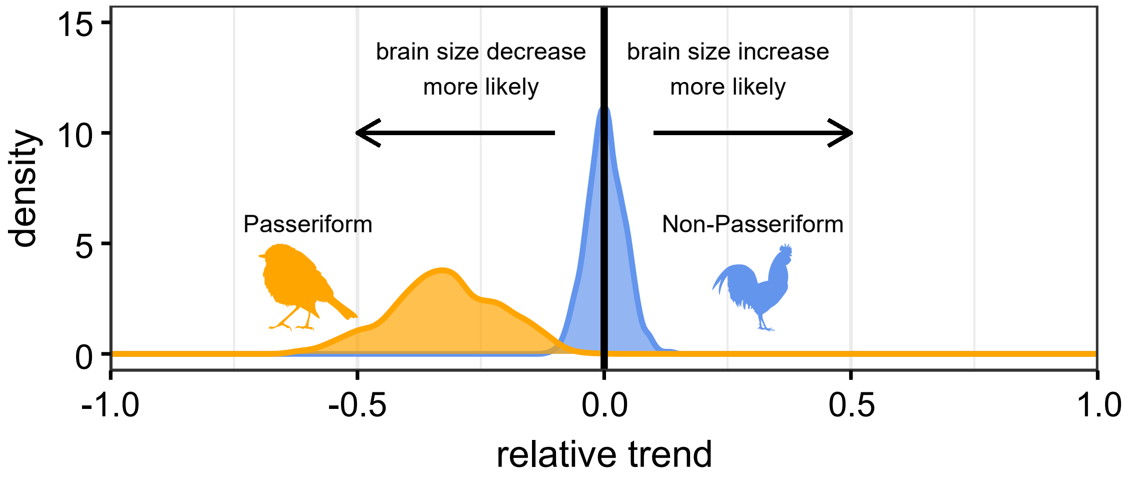


Fig 4: Estimated trends in relative brain size transitions on the 'early crown' tree (110 Ma). Values greater than zero indicate a bias toward brain size increases.

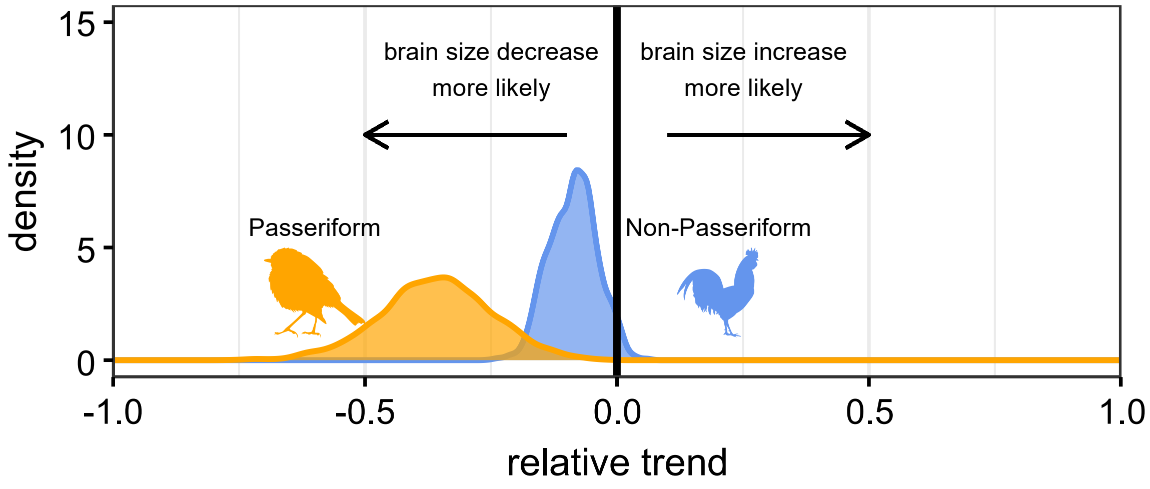


Fig 5: Estimated trends in relative brain size transitions on the 'middle crown' tree (92.51 Ma). Values greater than zero indicate a bias toward brain size increases.

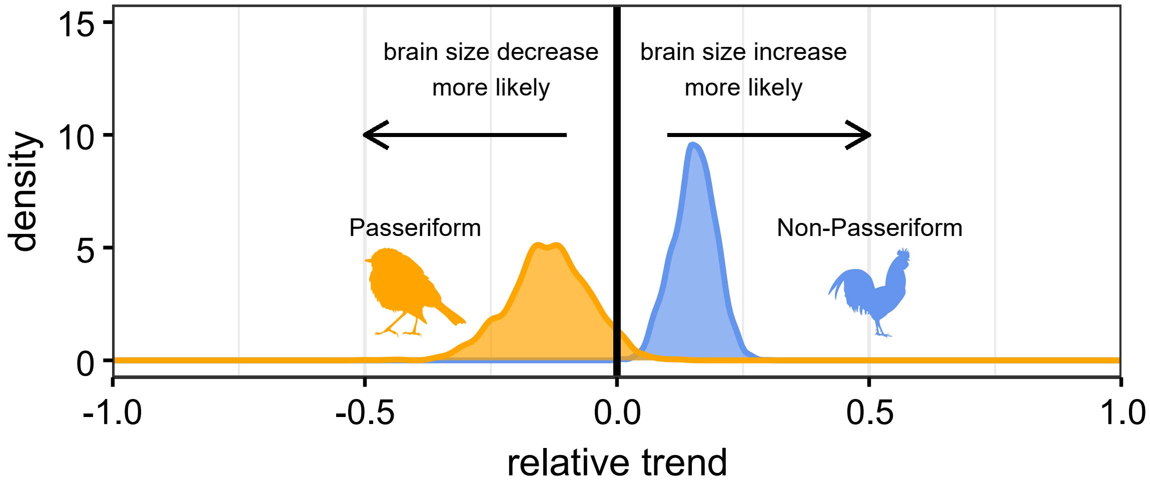


Fig 6: Estimated trends in relative brain size transitions on the 'late crown' tree (72 Ma). Values greater than zero indicate a bias toward brain size increases.

QuaSSE Model Results (Symmetric, No Bias Term)

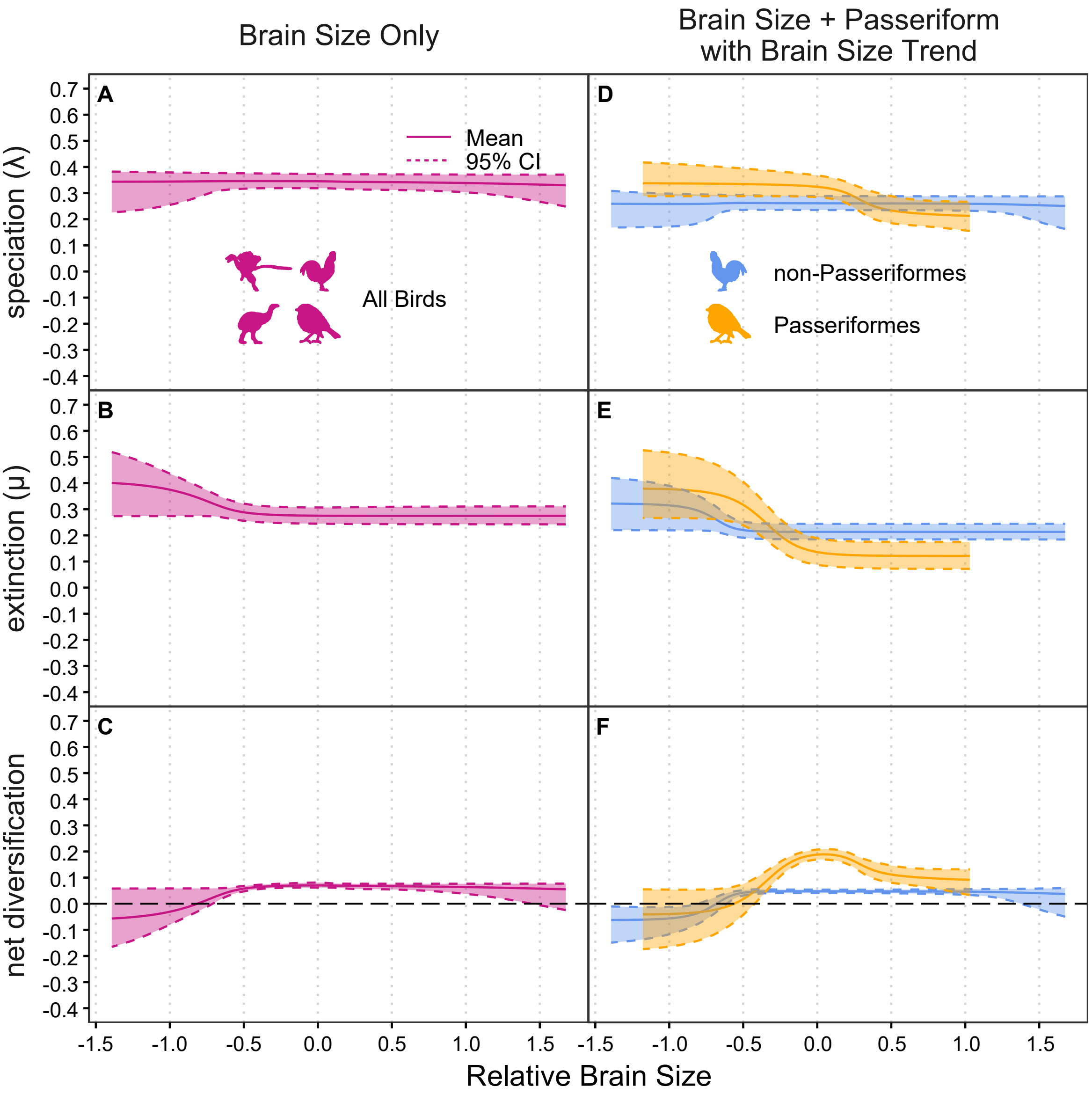


Fig 7: Diversification in birds (A-C) and for Passeriformes/non-Passeriformes (D-F) as a function of relative brain size for QuaSSE/BiQuaSSE models with symmetric rates of relative brain size increases and decreases. Residual brain size values from a linear regression against body size were used to control for the effect of allometry. Under the second model (D-F), the brain size transition rate is influenced by a bias term, allowing for trends in brain size evolution. Larger-brained lineages have lower speciation (A) and lower extinction (B) rates. Passeriformes have extinction rates lower than non-Passeriformes at highest brain sizes (E), leading to a peak in net diversification (F). Model parameters inferred using the "early crown" phylogeny (most recent common ancestor of crown birds at 110 Ma).

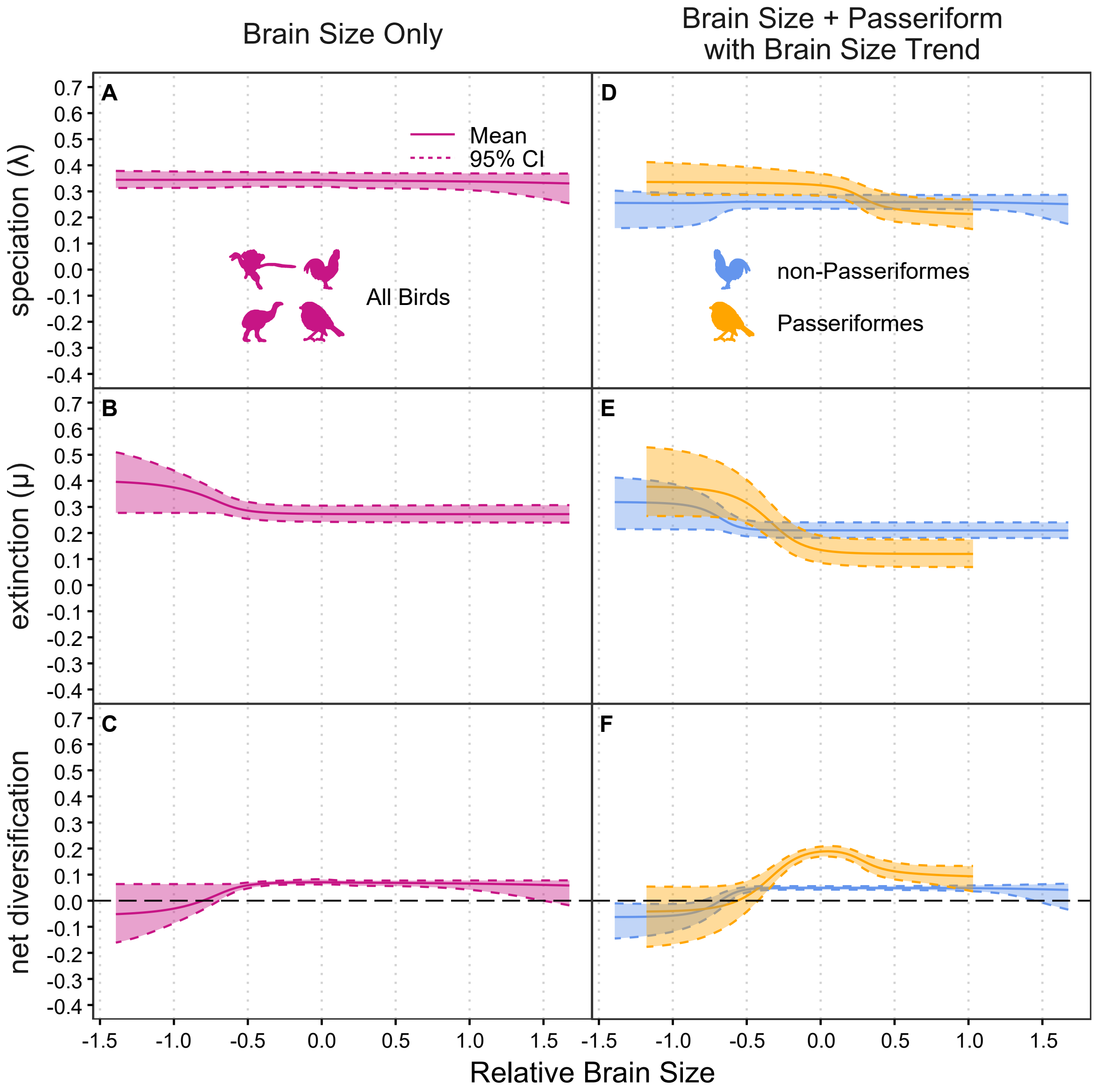


Fig 8: Diversification in birds (A-C) and for Passeriformes/non-Passeriformes (D-F) as a function of relative brain size for QuaSSE/BiQuaSSE models with symmetric rates of relative brain size increases and decreases. Residual brain size values from a linear regression against body size were used to control for the effect of allometry. Under the second model (D-F), the brain size transition rate is influenced by a bias term, allowing for trends in brain size evolution. Larger-brained lineages have lower speciation (A) and lower extinction (B) rates. Passeriformes have extinction rates lower than non-Passeriformes at highest brain sizes (E), leading to a peak in net diversification (F). Model parameters inferred using the "middle crown" phylogeny (most recent common ancestor of crown birds at 92.51 Ma).

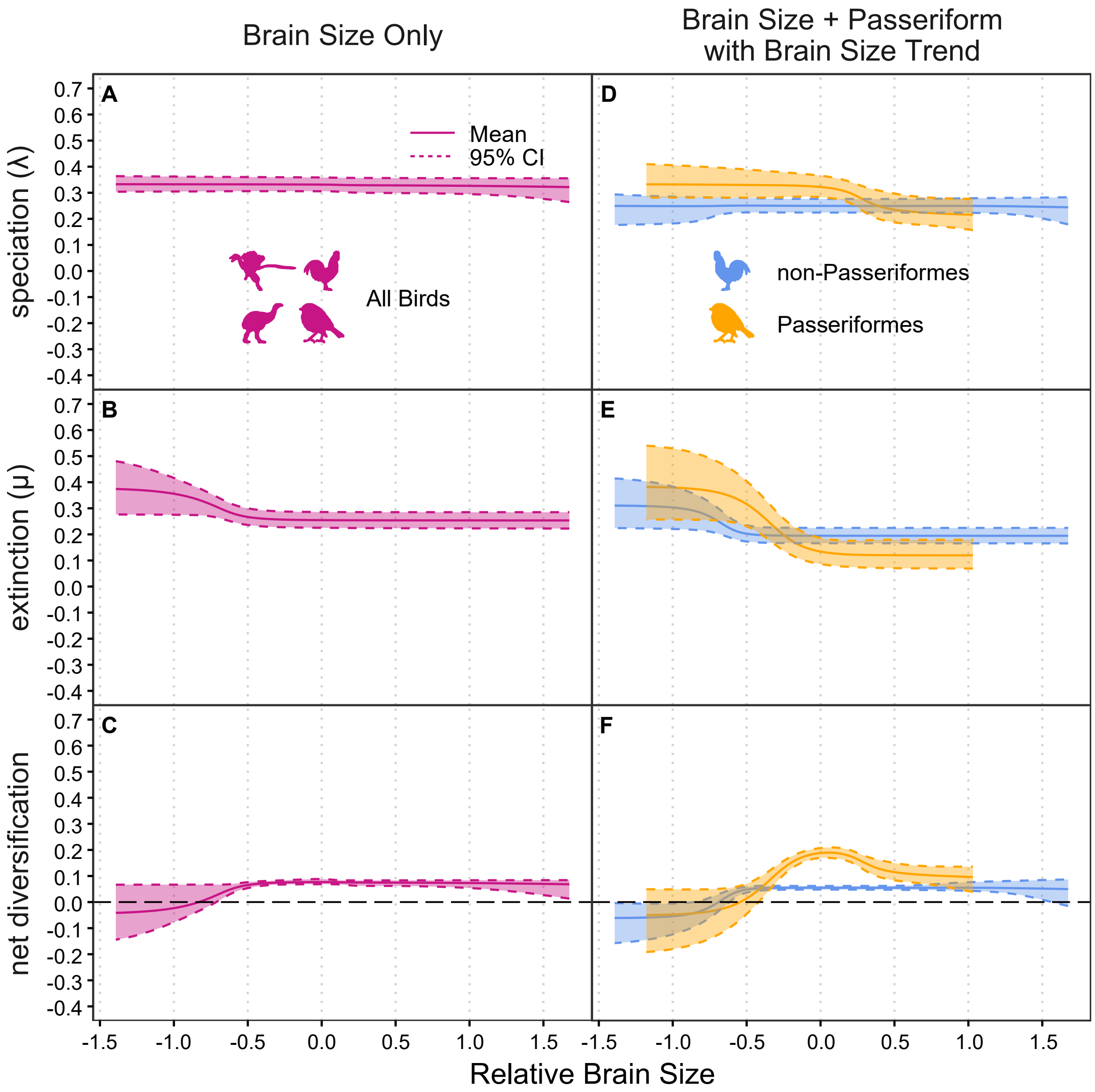


Fig 9: Diversification in birds (A-C) and for Passeriformes/non-Passeriformes (D-F) as a function of relative brain size for QuaSSE/BiQuaSSE models with symmetric rates of relative brain size increases and decreases. Residual brain size values from a linear regression against body size were used to control for the effect of allometry. Under the second model (D-F), the brain size transition rate is influenced by a bias term, allowing for trends in brain size evolution. Larger-brained lineages have lower speciation (A) and lower extinction (B) rates. Passeriformes have extinction rates lower than non-Passeriformes at highest brain sizes (E), leading to a peak in net diversification (F). Model parameters inferred using the "middle crown" phylogeny (most recent common ancestor of crown birds at 72 Ma).

QuaSSE Model Results (Extant Only)

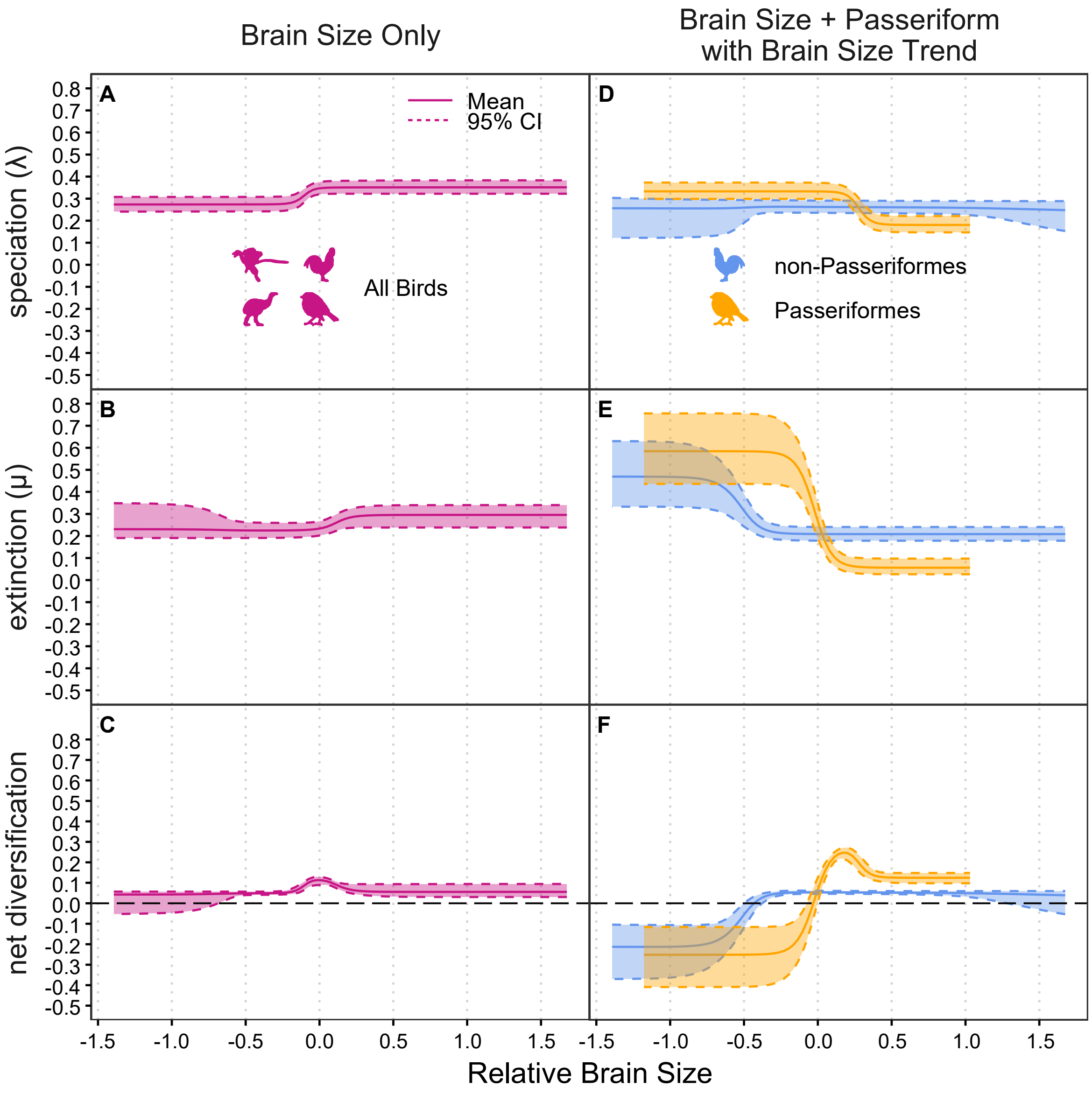


Fig 10: Diversification in birds (A-C) and for Passeriformes/non-Passeriformes (D-F) as a function of relative brain size in the extant only dataset. Residual brain size values from a linear regression against body size were used to control for the effect of allometry. Under the second model (D-F), the brain size transition rate is influenced by a bias term, allowing for trends in brain size evolution. Larger-brained lineages have lower speciation (A) and lower extinction (B) rates. Passeriformes have extinction rates lower than non-Passeriformes at highest brain sizes (E), leading to a peak in net diversification (F). Model parameters inferred using the “early crown” phylogeny (most recent common ancestor of crown birds at 110 Ma).

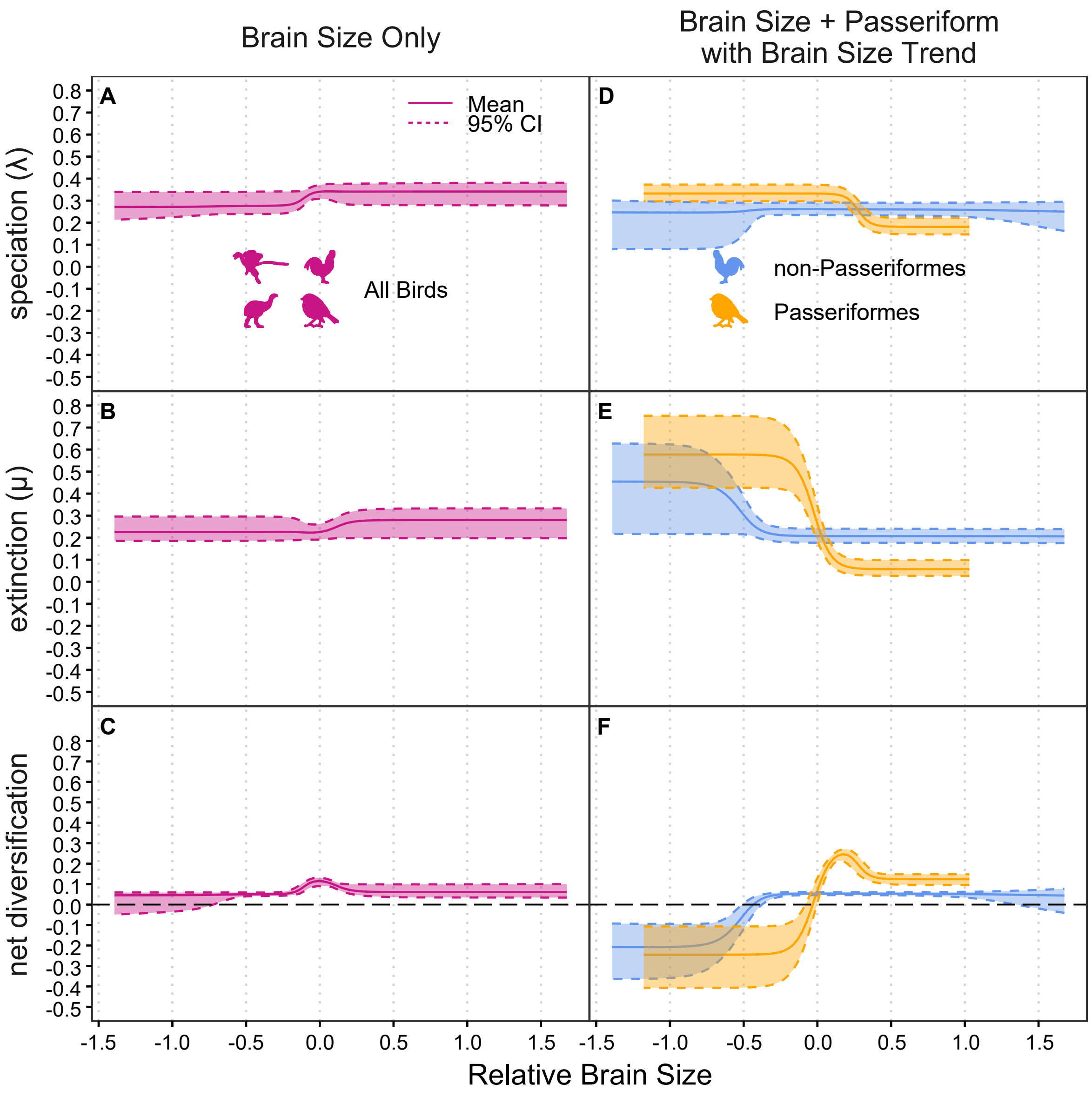


Fig 11: Diversification in birds (A-C) and for Passeriformes/non-Passeriformes (D-F) as a function of relative brain size for the extant only dataset. Residual brain size values from a linear regression against body size were used to control for the effect of allometry. Under the second model (D-F), the brain size transition rate is influenced by a bias term, allowing for trends in brain size evolution. Larger-brained lineages have lower speciation (A) and lower extinction (B) rates. Passeriformes have extinction rates lower than non-Passeriformes at highest brain sizes (E), leading to a peak in net diversification (F). Model parameters inferred using the "middle crown" phylogeny (most recent common ancestor of crown birds at 92.51 Ma).

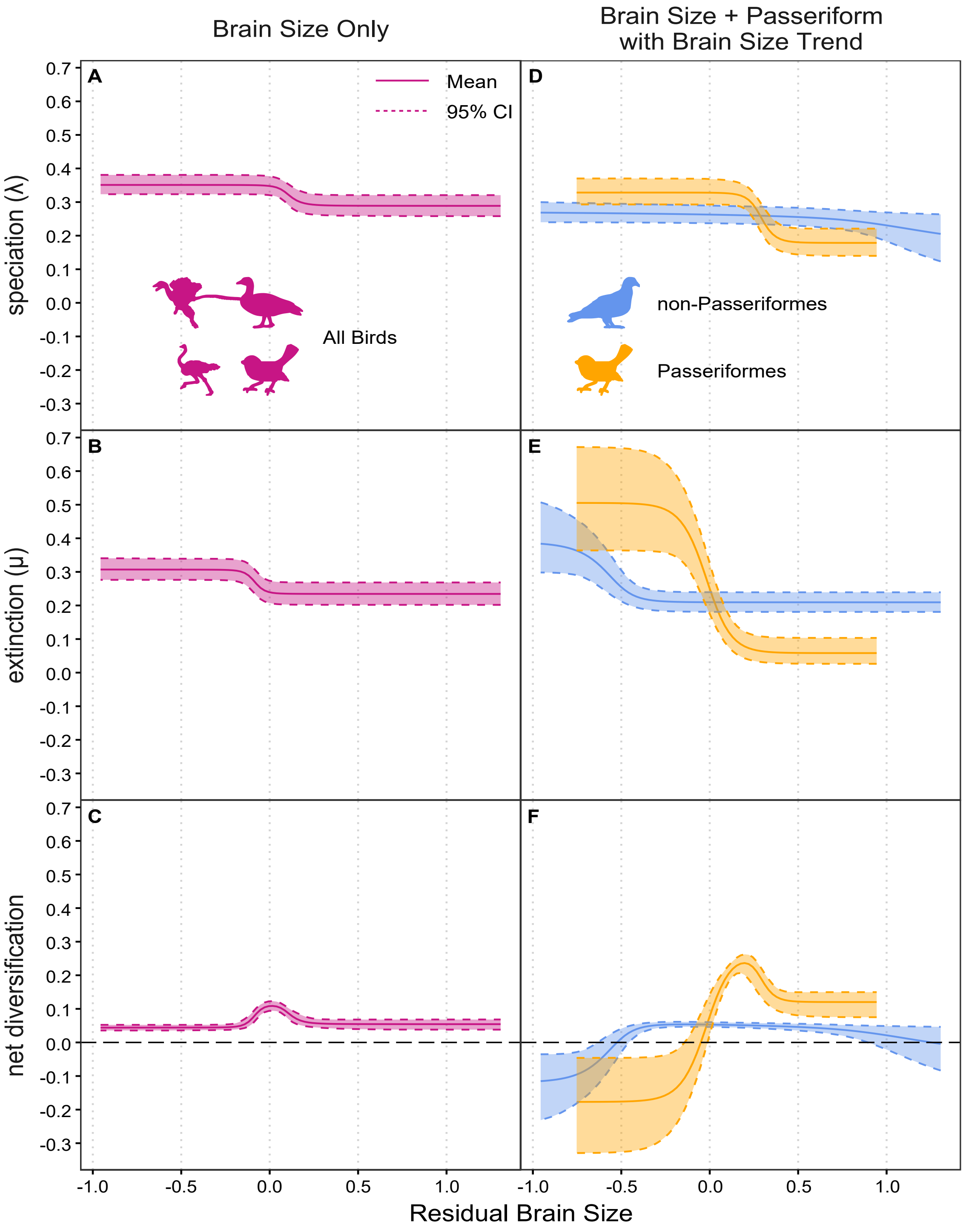


Fig 12: Diversification in birds (A-C) and for Passeriformes/non-Passeriformes (D-F) as a function of relative brain size for the extant only dataset. Residual brain size values from a linear regression against body size were used to control for the effect of allometry. Under the second model (D-F), the brain size transition rate is influenced by a bias term, allowing for trends in brain size evolution. Larger-brained lineages have lower speciation (A) and lower extinction (B) rates. Passeriformes have extinction rates lower than non-Passeriformes at highest brain sizes (E), leading to a peak in net diversification (F). Model parameters inferred using the “late crown” phylogeny (most recent common ancestor of crown birds at 72 Ma).

**Table 1: Tip-rate correlations from pgls tests**

| Dataset | Crown divergence time | Adjusted $R^2$ | p-value |
|---|---|---|---|
| All taxa (extant & fossil) | Early (110 M.y.a.) | 0.002093 | 0.01607 |
| | Middle (92.5 M.y.a) | 0.002097 | 0.016 |
| | Late (73 M.y.a) | 0.001774 | 0.02444 |
| Extant only | Early (110 M.y.a.) | 0.00224 | 0.01381 |
| | Middle (92.5 M.y.a) | 0.002417 | 0.01102 |
| | Late (73 M.y.a) | 0.002276 | 0.01318 |
| Passeriformes (perching birds) only | Early (110 M.y.a.) | -0.0001778 | 0.3631 |
| | Middle (92.5 M.y.a) | -0.0001778 | 0.3631 |
| | Late (73 M.y.a) | -0.0001778 | 0.3631 |
| Non-Passeriformes only | Early (110 M.y.a.) | 0.002177 | 0.04903 |
| | Middle (92.5 M.y.a) | 0.002414 | 0.04069 |
| | Late (73 M.y.a) | 0.001959 | 0.05825 |

**Table 2: Tip-rate correlations from ES-sim tests**

| Dataset | Crown divergence time | Pearson's correlation coefficient (ρ) | p-value |
|---|---|---|---|
| All taxa (extant & fossil) | Early (110 M.y.a.) | -0.04337821 | 0.7272727 |
| | Middle (92.5 M.y.a) | -0.04129063 | 0.6953047 |
| | Late (73 M.y.a) | -0.03567743 | 0.6993007 |
| Extant only | Early (110 M.y.a.) | -0.02347432 | 0.8171828 |
| | Middle (92.5 M.y.a) | -0.02107855 | 0.8751249 |
| | Late (73 M.y.a) | -0.01553814 | 0.9110889 |
| Passeriformes (perching birds) only | Early (110 M.y.a.) | 0.02996528 | 0.8251748 |
| | Middle (92.5 M.y.a) | 0.02996528 | 0.8011988 |
| | Late (73 M.y.a) | 0.02996528 | 0.8431568 |
| Non-Passeriformes only | Early (110 M.y.a.) | 0.0513571 | 0.6813187 |
| | Middle (92.5 M.y.a) | 0.0522314 | 0.6953047 |
| | Late (73 M.y.a) | 0.05011265 | 0.6593407 |
